# Developability Evaluation of Single-Domain Antibody Chelator Conjugates for Diagnostic Radiotracers

**DOI:** 10.64898/2026.02.09.704800

**Authors:** Philipp D. Kaiser, Simon Straß, Sandra Maier, Evgenia Herbold, Bjoern Traenkle, Anne Zeck

**Affiliations:** NMI Natural and Medical Sciences Institute at the University of Tübingen, Reutlingen, Germany

**Keywords:** Single-domain antibodies (sdAbs), developability assessment, chelator conjugates (NODAGA), Chelatability (Cu^2+^ radiolabeling efficiency), Dynamic light scattering (DLS), Liquid chromatography-mass spectrometry (LC–MS), nanoDSF (thermal stability), Peptide mapping (LC–MS/MS), quantitative Antigen Binding Capacity SEC (qABC-SEC)

## Abstract

**Background/Objectives:** Developability assessment is a critical step in advancing antibody-based molecules toward clinical application. This evaluation typically begins during clinical candidate selection and continues throughout all modifications of the molecule during development. It is guided by the target product profile, which includes the intended administration route and regimen, formulation parameters, and process conditions encountered during manufacturing, storage, and delivery. While developability testing is well established for conventional therapeutic antibodies, strategies for assessing single-domain antibodies (sdAbs) and their conjugates remain underexplored. Here we present a strategy to test the developability of sdAbs as a case study for two clinical candidates intended as precursors for the production of diagnostic tracers for clinical imaging.

**Methods:** Assays were developed to evaluate chemical and thermodynamic stability, target binding affinity and capacity, and chelation efficiency (“chelatability”). Accelerated stability studies were conducted for both unconjugated sdAbs and their chelator conjugated forms following incubation at two pH conditions, at multiple time points, and after twelve freeze-thaw cycles to simulate process conditions and long-term storage. Analytical assays were applied stepwise in a hierarchical approach to minimized experimental effort and material consumption. Candidates exhibiting critical developability features were selectively addressed by assays with increasing precision.

**Results:** A tailored panel of analytical assays optimized for low molecular weight proteins was established and applied to the two clinical candidates, identifying instability hotspots as well as potential mitigation strategies. Successful engineering of a candidate with an initially critical developability profile was achieved.

**Conclusion:** This study demonstrates the implementation of a structured developability assessment strategy for sdAb conjugates. The approach integrates physicochemical and functional stability evaluations, supporting robust candidate selection, formulation development, and method optimization for this class of molecules.

## 1. Introduction

In the last 40 years, since the first monoclonal antibody drug [1], more than 200 antibody-based therapeutics have entered the clinic across diverse indications [2]. While still dominated by conventional IgG molecules, the clinical antibody field has expanded to engineered formats such as antibody fragments, bispecific antibodies, and antibody-drug conjugates (ADCs) [3]. For therapeutic and diagnostic scenarios, where improved tissue penetration, target accessibility, and short half-life is needed, various approaches of antibody miniaturization have been explored [4,5].

In recent years, single-domain antibodies (sdAbs), the variable domains derived from camelid heavy-chain antibodies, have gained increasing attention as an emerging class of biotherapeutics and diagnostic agents [6]. Caplacizumab, the first sdAb-based therapeutic, received FDA and EMA approval in 2018 for the treatment of acquired thrombotic thrombocytopenic purpura, marking the clinical debut of this molecular class and paving the way for numerous preclinical and clinical sdAb programs in oncology, inflammation, and infectious disease [7].

Beyond their therapeutic potential, sdAbs are particularly well suited for molecular imaging applications, including positron emission tomography (PET) [8]. Their small size (∼15 kDa), entails rapid blood clearance and high tissue penetration, making them ideal candidates for targeted imaging. Several sdAb-based tracers have advanced to clinical evaluation, such as ^68^Ga-NOTA-anti-HER2 sdAbs for breast cancer imaging and ^68^Ga-NOTA-anti-MMR sdAbs for macrophage imaging in oncology [9,10]. These examples highlight the growing role of sdAbs as diagnostic tracers for patient stratification and therapy monitoring, complementing the established field of therapeutic antibodies. A recent advance supporting the rapid development of sdAb technologies is the generation of highly diverse and stable synthetic libraries, which enable efficient selection of sdAbs against even challenging targets, as demonstrated by Zimmermann et al. [11].

This rapid growth of therapeutic IgG antibody pipelines has driven the establishment of dedicated developability assessment frameworks to mitigate risks in antibody drug development. These frameworks aim to identify molecular liabilities early and to de-risk candidates prior to large-scale manufacturing and clinical evaluation [12]. Typical developability assessments encompass chemical and physical stability, aggregation propensity, immunogenicity risk, expression yield, and formulation compatibility [13]. To support these efforts, a wide range of analytical methods have been developed and standardized. Collectively, these tools have enabled the successful translation of antibody-based therapeutics into clinically and commercially viable products.

However, developability and analytical assessment frameworks developed for conventional antibodies are not readily transferable to sdAbs and their conjugates. Their smaller size, distinct surface charge distribution, and lack of an Fc region influence parameters such as thermal stability, solubility, and aggregation behavior, and may also alter interactions with excipients and container materials. Furthermore, when sdAbs are chemically modified, for example, by conjugation with metal chelators, additional factors such as chelate integrity and metal-binding efficiency become critical for product quality and functional performance [14]. Consequently, adapted developability strategies and specialized analytical methods are required to ensure that sdAb conjugates maintain stability, binding activity, and chelatability throughout manufacturing, storage, and clinical use.

While others have reported on approaches to isolate sdAbs with favorable developability [15] or to predict developability in silico [16], little has been published on experimental developability assessment of clinical sdAb candidates [17]. The approach presented here integrates physicochemical and functional stability assessment, providing a structured framework for robust candidate selection and formulation development of sdAb-tracer conjugates intended for diagnostic imaging applications.

## 2. Materials and Methods

### Proteins and Chemicals

All chemicals were obtained from commercial suppliers and used without further purification. Unless otherwise stated, reagents were of analytical grade or higher. The chelator p-NCS-benzyl-NODAGA was purchased from CheMatech (Dijon, France). Recombinant human SIRPα (Antigen 2) was acquired from ACROBiosystems and used for affinity measurements via biolayer interferometry. Single-domain antibodies (sdAb1, sdAb2 wt and variants) and the antigen 1-Fab fusion protein (huCD4___domain___1&2-Fab), used for antigen-binding analysis were produced in-house. Expression was performed in ExpiCHO cells via transient transfection with pcDNA3-based plasmids. Proteins were purified from cell-free culture supernatants by affinity chromatography using HiTrap™ MabSelect™ VH3 columns, followed by desalting with HiPrep™ 26/10 Desalting columns (Cytiva, Marlborough, MA, USA).

### Preparation of sdAb Chelator Conjugates

Conjugation of sdAbs with NODAGA was achieved using a site-selective amine-reactive strategy, yielding a well-defined and homogeneous conjugate. Briefly, a twelve-fold molar excess of the NCS reagent dissolved in water was added to purified sdAbs in coupling buffer (0.2 M potassium phosphate, pH 8.0). The reaction mixture was incubated for 48 hours at 35 °C. Excess chelator was removed by size exclusion chromatography using a HiLoad™ Superdex 75 pg column (Cytiva) equilibrated with PBS.

### Preparation of Stability Samples

SdAbs and sdAb conjugates were buffer-exchanged from the initial formulation buffer (20 mM sodium phosphate pH 6.8) into 20 mM His HCl buffer, 140 mM NaCl pH 6 and 50 mM TRIS HCl buffer pH 8, respectively. Briefly, 200-400 µL (corresponding to 1.2 mg protein) were buffer-exchanged using Amicon® Ultra-4 filters (Merck Millipore) with a molecular weight cut-off of 3 kDa as described in the technical manual. Samples were diluted to a final volume of 3000 µL in the corresponding stress buffer, mixed and centrifuged for 30 minutes at 7500 x g to obtain ∼300 µL retentate. This process was repeated once. Concentration was adjusted to 1.5 mg/mL for all samples in the corresponding stress buffer and determined by NanoDrop spectrophotometer (Thermo Fisher Scientific) using theoretically calculated extinction coefficients by protpi (https://www.protpi.ch/).

Buffer-exchanged stability samples were incubated at 40 °C for one and four weeks, respectively to assess potential degradation hot spots. In addition, sdAb conjugates in initial buffer composition were stored at 4 °C for one and four weeks to assess stability of conjugation. As reference samples, buffer-exchanged and original samples were stored frozen at -80 °C. After the respective stress duration, the samples were immediately frozen at -80 °C until sample preparation and analysis.

Freeze/thaw experiments were conducted by freezing 50 µL aliquots of sdAb1 (7.4 mg/mL) and sdAb1-NODAGA (2.85 mg/mL), respectively in phosphate buffer pH 8 for 15 to 30 minutes at -8 to -10 °C followed by thawing the samples by placing them for 10 minutes on a ThermoMixer® C (Eppendorf, Germany) at 4 °C. The cycle was repeated 12 times.

### Intact Mass Analysis for Integrity Analysis

1.5 µg (∼1 µL) of sdAb and sdAb-conjugate samples were analyzed by LC-MS using an UHPLC system (UltiMate 3000, RSLCnano, Dionex GmbH, Idstein, Germany) coupled to a QTOF-type mass spectrometer (maXis II QTOF, Bruker, Bremen, Germany). Reversed-phase chromatography (ACQUITY BEH C4, 1.7 µm, 300 Å, 1 mm × 50 mm, Waters GmbH, Eschborn, Germany) was applied for desalting. The solvents were 0.1 % formic acid in (A) water and in (B) acetonitrile, respectively. A stepwise linear gradient was run at 75 °C at a flow rate of 150 µL/min (0 - 0.4 min, 5 % B; 2.5 min 30 % B; 7 min, 50%B) followed by wash steps. Mass spectrometer parameters were adapted to the size of the molecule and the flow rate. The first two minutes of the LC eluate were directed into waste using the valve of the QTOF. External calibration was performed using ESI Low Concentration Tuning Mix (Agilent Technologies) and recalibration of each dataset was performed as part of a postprocessing method using tuning mix inserted during a calibration segment (run time 2 - 2.1 minutes). Data analysis was performed by summing up the mass spectra in the relevant retention time window followed by charge-deconvolution using the MaxEnt algorithm provided by the Bruker Compass DataAnalysis software version 6.1. Assignment of mass peaks was achieved by matching theoretical and experimental mass.

### Biolayer Interferometry (BLI) for Affinity Evaluation of sdAb2 Wt and Variants

Equilibrium constant, K_D_, the association rate constant, K_a_, and the dissociation rate constant, K_d_, of antigen 2 and sdAb2 wt or variant was determined by biolayer interferometry using Octet RED96e system (Sartorius, Göttingen, Germany) as per the manufacturer’s recommendations. In brief, biotinylated hSIRPα (5 µg/mL) diluted in Octet buffer (PBS, 0.1 % BSA, and 0.02 % Tween-20) was immobilized on streptavidin-coated biosensor tips (SA, Sartorius) for 120 seconds. In the association step, a dilution series of sdAb2 or sdAb2-NODAGA ranging from 0.625 nM to 320 nM were reacted for 240 s followed by dissociation in Octet buffer for 720 s. Every run was normalized to a reference run applying Octet buffer for association. Data were analyzed using the Octet Data Analysis HT 12.0 software applying the 1:1 ligand-binding model and global fitting.

### Size Exclusion Chromatography (SEC) for sdAb Antigen Binding Capacity

SEC was performed to assess the binding capacity of sdAb1 samples (reference and stability) to the target antigen fused to a non-interacting Fab. For preparation of the sdAb-antigen mixtures, a total of 3.6 µM sdAb1 material (reference and stability) was mixed with 7.2 µM of target antigen in DPBS buffer and incubated at RT in PCR tubes for 10 minutes. Samples were then centrifuged and transferred to HPLC vials. Then, 20 µL of the mixture was injected into the HPLC for SEC separation.

The analysis was conducted using an Agilent 1260 Infinity HPLC system, equipped with a variable wavelength detector (VWD) recording absorption at 220 nm. Chromatographic separation was achieved using a TSKgel SuperSW2000 column (4.6 × 300 mm, 4 µm, 125 Å silica, Tosoh Bioscience), preceded by a TSKgel SuperSW Guard-column (4.6 × 35 mm, 4 µm, Tosoh Bioscience). The mobile phase consisted of a phosphate buffer (pH 7.0), containing K_2_HPO_4_ 140 mM, KH_2_PO_4_ 60 mM and KCl 250 mM. The flow rate was set to 0.25 mL/min, with a column temperature of 25 °C. An injection volume of 20 µL was used, and the total run time per sample was 30 minutes.

### Tryptic Peptide Map Analysis

#### Tryptic Digestion Protocol at pH 7.5

The stressed and reference samples were denatured and reduced by mixing 50 µg protein sample (33 µL per sample) with denaturing buffer to give a final volume of 108 µL (0.4 M Tris/HCl, 8.0 M Gua-HCl, pH 8.0) and 4 µL of 0.24 M freshly prepared DTT solution in denaturing buffer. The reduction was carried out at 37 °C for 1 hour (550 rpm). The reduced samples were cooled to room temperature and subsequently alkylated by addition of 4 µL freshly prepared alkylation reagent (0.6 M iodoacetamide in water). The alkylation process was carried out at room temperature in the dark for 15 minutes. The excess of alkylation reagent was inactivated by addition of 4 µL of DTT solution. The samples were then buffer-exchanged to approx. 120 µL of 50 mM Tris/HCl, 100 mM methionine, pH 7.5 using Zeba… Spin desalting column (Thermo Fisher Scientific) by adding three times new buffer and centrifuging at 1500 g for 1 minute. Digestion was performed with mass spectrometry-grade trypsin for 20 h at 37 °C at an enzyme to substrate ratio of 1:50 (w/w) by addition of 4 µL reconstituted trypsin solution. The digestion was stopped by addition of 4 µl 10 % trifluoroacetic acid.

#### Tryptic Digestion Protocol at pH 6.0 for Assessment of Succinimide Formation

The samples 1 - 4 were denatured and reduced by mixing 50 µg of sample (33 µL per sample, respectively) with denaturing buffer to give a final volume of 116 µL (0.2 M His/NaCl, 8.0 M Gua-HCl, pH 6.0) and 4 µL of freshly prepared 500 mM TCEP solution in denaturing buffer. The reduction was carried out at 37 °C for 1 hour. The reduced sample was cooled to room temperature. No alkylation was performed. The samples were then buffer-exchanged to approx. 120 µL of 0.2 M His/NaCl, 0.5 mM TCEP-HCl pH 6.0 as described above. Digestion was performed with trypsin for 20 h at 37 °C at an enzyme to substrate ratio of 1:50 (w/w). The digestion was stopped by addition of 4 µl 10 % trifluoroacetic acid.

### On-line Reversed-Phase Chromatography Tandem Mass Spectrometry (LC-MS/MS)

The samples were injected and separated using reversed-phase HPLC (Vanquish Horizon, Thermo Fisher Scientific). A capLC column (Waters, Acquity Peptide CSH C18, 130 Å, 1.7 µm, 1 mmx150mm) was used for separation of the peptides. A 90-minute linear gradient was applied as follows (minute/%B): 0/2.5, 3/2.5, 33/50, 34/99, 38/99, 40/2.5, 60/2.5. Eluent A was H_2_O with 0.1 % formic acid and eluent B was 80 % acetonitrile with 0.1 % formic acid. The injection amount was 10 µL. The flow rate was 100 µL/min, the column oven temperature was 65 °C. The HPLC eluate was directly infused into an Eclipse mass spectrometer (Thermo Fisher Scientific). The mass spectrometer operated in positive ion mode, the spray voltage was 3.5 kV, the capillary temperature was 250 °C and vaporizer temperature 75 °C. For the MS/MS product ion scan, the activation type was higher energy CID (HCD). The 4-scan-event method applied consisted of a full MS survey scan at m/z 200-2000 and resolution power (RP) of 120 000 followed by three cycles of data-dependent MS/MS scans on the top three most intense ions. The dynamic exclusion function was enabled and parameters were as follows: dynamic exclusion of 90 sec, isolation window is set automatically, resolution of 30 000, normalized AGC target 2000 % (MS/MS), maximum inject time Auto, HCD collision energy of 25 %, intensity threshold of 5e4. Unassigned charge states and charge state +1 and ≥8 were rejected for MS/MS triggering.

For data processing, the Mascot 3.0 error tolerant search (Matrix Science London, UK) based on an in-house protein database that contained the sequences of the sdAbs. MS/MS spectra were charge-deconvoluted using MS2 processor before database search and Mascot settings were adapted to allow identification of small peptides: the shortest peptide tested in the search and evidenced in the report was set to 4 amino acids. Mascot ETS search parameters were: enzyme was trypsin, 1 missed cleavage allowed, fixed modification carbamidomethylation for pH 7.5 digest, peptide tolerance: 6 ppm, MS/MS tolerance 0.05 Da, peptide charge +2, +3, +4, error tolerant search activated, and confident peptide identification. Additionally, modifications were identified and relatively quantified using the software BiopharmaFinder 4.1 (Thermo Scientific).

### NanoDSF and DLS Assays

Thermal stability and dynamic light scattering assays. The dynamic light scattering and thermal stability of sdAb1, sdAb1-NODAGA and sdAb1-NODAGA with Cu^2+^ was monitored (NanoTemper Technologies) at a concentration of 1 mg/ml in 20 mM sodium phosphate, pH 6.8 buffer. Standard grade nanoDSF capillaries (NanoTemper Technologies) were loaded into a Prometheus Panta device (NanoTemper Technologies) controlled by PR.PantaControl (version 1.8). Isothermal measurements for DLS were collected at 25 °C using the high-sensitivity mode prior to the thermal denaturation experiments. For nanoDSF, excitation power was adjusted to 30 % and samples were heated from 20 °C to 90 °C with a slope of 1 °C/min. All samples were run in triplicates and errors represent standard deviations. The stability of the samples was analyzed in independent experiments.

## 3. Results

### 3.1. Design of Developability Assays for sdAb and sdAb-Chelator Conjugates

The case study focuses on single-domain antibodies (sdAbs) and their chelator conjugates intended for radiolabeling and clinical PET imaging. SdAbs are produced recombinantly and may undergo interim storage prior to conjugation with the NODAGA chelator. As precursors for the final radiolabeling step, sdAb-chelator conjugates must retain longterm stability, chelatability, and structural integrity during manufacturing, storage, and shipment as injection-ready radiopharmaceuticals (Figure 1). With proof-of-function for expression, purification, and radiolabeling already established, this study primarily evaluates molecular liabilities relevant for long-term storage of both unconjugated sdAbs and their conjugated forms.

**Figure 1:**
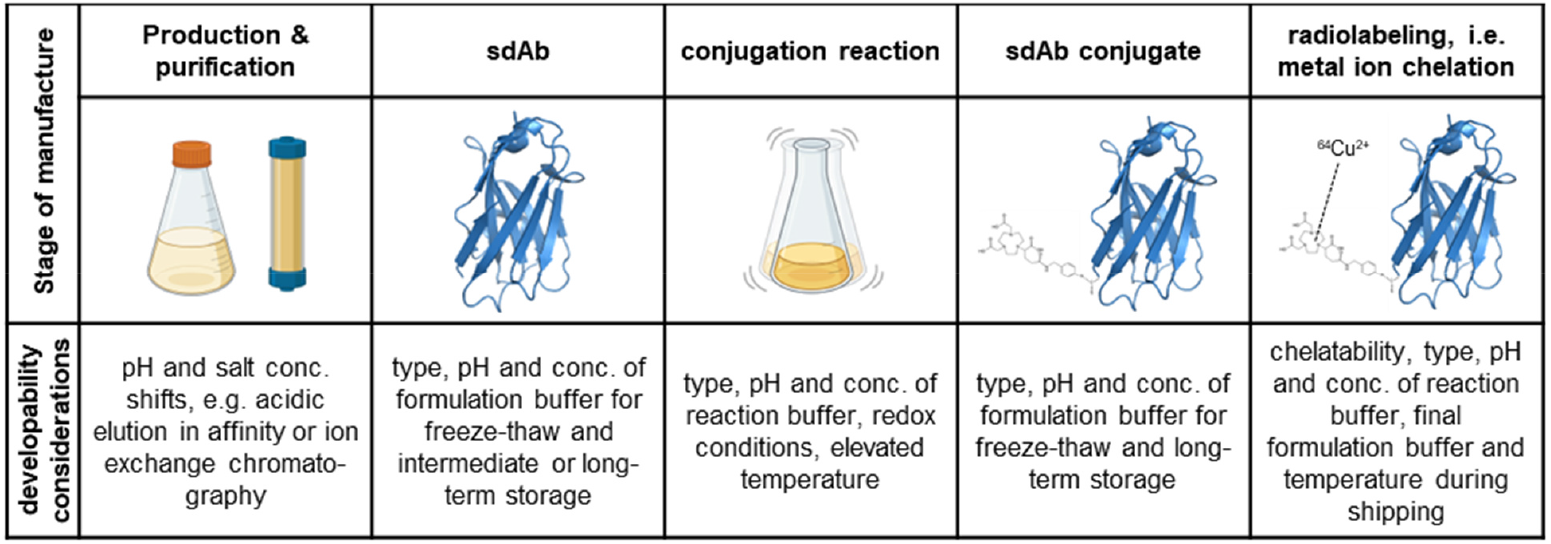
Stages of manufacturing of sdAb-chelator conjugates. Figure created with BioRender.com.

To address the relevant process and storage conditions, a set of targeted stress experiments was designed. SdAbs and sdAb-chelator conjugates were exposed to acidic and basic pH, elevated temperature, and freeze-thaw cycling to simulate potential degradation scenarios. The stressed samples were analyzed for: non-enzymatic proteolysis via intact-mass LC-MS, chemical degradation (deamidation, succinimide formation, oxidation) via tryptic peptide mapping (LC-MS/MS), loss of antigen binding via SEC-based complex formation assays, alterations in tertiary structure via nanoDSF and DLS, chelator integrity and metal-binding efficiency via a mass-spectrometry-based chelation assay across different buffer systems. An overview of stability conditions and analytical methods applied is provided in *Table 1*.

**Table 1.** Design of stability-predicting assays for sdAb chelator conjugates.

| Stability parameter | Basic pH condition<br>(Tris-HCl, pH 8.0, 40 °C; 1 & 4 weeks) | Acidic pH condition<br>(His-NaCl, pH 6.0, 40 °C; 1 & 4 weeks) | Freeze-thaw stress<br>(12 cycles; phosphate buffer, pH 7.5) |
| --- | --- | --- | --- |
| CDR region stability | Asn/Asp degradation (deamidation, succinimide formation) |  | Not tested |
| Framework integrity | Non-enzymatic proteolysis; N-terminal pyroglutamate formation |  | Not tested |
| Structural integrity | NanoDSF (unfolding temperature, F350/330 ratio); DLS (PDI) |  |  |
| Antigen-binding capacity | SEC-UV complex-shift assay |  | Not tested |
| Chelatability (metal binding) | LC-MS-based chelation assay in multiple storage buffers |  | Not tested |

**Table 2.** Potential degradation hot spots in investigated sdAbs. Numbering system used is according to Abhinandan and Martin [24].

| Molecule region | sdAb1 | sdAb2 |
| --- | --- | --- |
| CDR1 | -- | -- |
| CDR2 | D52-S52a | D52b-G53, N56-T57, D61-S62 |
| CDR3 | D97-P98 | D100f-P100g |

### 3.2. In silico Prediction of Chemical Stability

Numerous studies describe computational approaches for predicting so-called *degradation hotspots* in proteins [18]. For example, asparagine deamidation is frequently observed at NG, NS, and NT sequence motifs, particularly when these residues are located in solvent-exposed and conformationally flexible structural elements. Likewise, succinimide formation and aspartate isomerization occur predominantly at DG and DS sequence motifs, while DP represents a known clipping hotspot [19-21]. Chemical degradation pathways and their impact on protein function and stability have been extensively characterized in the framework regions of antibodies [22,23]. In contrast, experimental validation is typically still required for modifications occurring in the complementarity-determining regions (CDRs). Consequently, stability assessments during developability studies usually focus on degradation events in the CDRs, while degradation observed in the framework regions often serves as a positive control in stress experiments.

When the N-terminal amino acid is used for chelator coupling, its intrinsic stability - particularly with respect to pyroglutamate formation - must also be evaluated. In the case of single-domain antibody-chelator (sdAb-chelator) conjugates, an additional degradation pathway must be considered: the loss of the conjugated molecule via bond hydrolysis in the N-terminal region. The susceptibility to such degradation depends on the steric properties of the conjugated moiety, as well as the length, chemistry, and configuration of the linker and conjugation bond. Taking these aspects into account, the following potential degradation hotspots were identified in the investigated molecules.

### 3.3. Assessment of Degradation Liabilities in the sdAb CDR Regions

#### Succinimide formation

can occur from aspartate and asparagine residues at pH 6 and above, particularly when they are followed by a small hydrophilic residue in the primary sequence. To evaluate whether such a reaction takes place in the investigated sdAbs, we first assessed the degree of succinimide formation via intact-mass LC-MS analysis. To differentiate succinimide-related loss of water from the loss of water associated with N-terminal pyroglutamate formation, we examined the intact-mass profiles of the sdAb-chelator conjugates. As shown in Figure 2A and 2C, sdAb1-NODAGA exhibits only minimal increases in water or ammonia loss at the intact-molecule level, indicating that D52-S52a does not undergo succinimide formation under the applied conditions. In contrast, sdAb2-NODAGA displays a marked increase in water (or ammonia) loss at the intact-molecule level already after one week of incubation at pH 6 and 37 °C, as illustrated in Figure 2B and 2D. This observation is consistent with significant succinimide formation occurring in sdAb2-NODAGA under these conditions.

**Figure 2.**
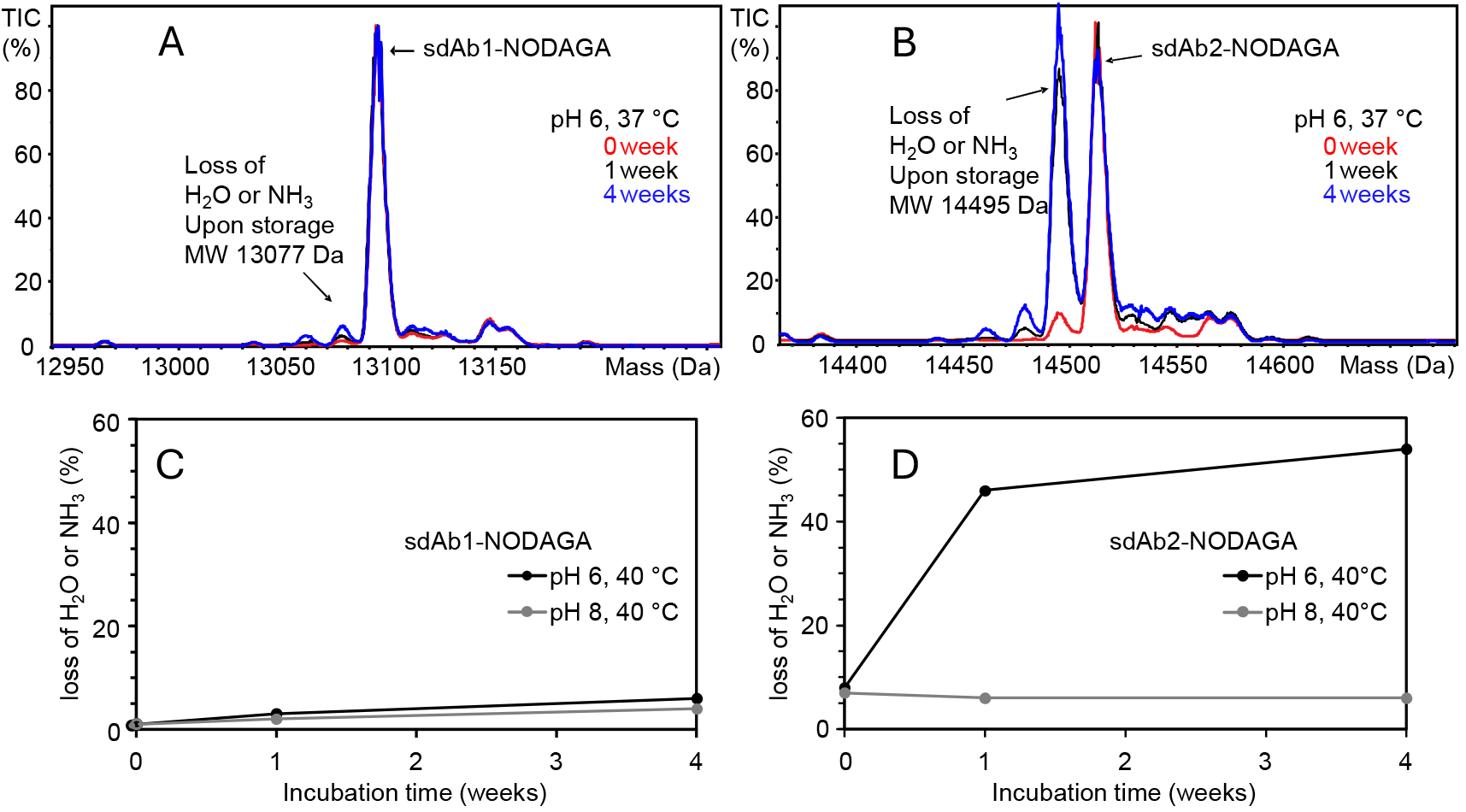
Assessment of succinimide formation from Asp or Asn by intact mass RP-C4-LC-MS analysis of sdAb-chelator samples stored at pH 6 and 8 at 37 °C. A, B: charge-deconvoluted mass spectra of sdAb conjugates after storage at pH 6 and C, D relative quantification of loss of water (or ammonia) using peak intensities of charge-deconvoluted mass spectra.

The loss of water (or ammonia) in sdAb2 was further investigated at peptide level to assign the degradation hot spot. In order to preserve the succinimide during sample preparation, pH 6 was used for tryptic digestion even though this sub-optimal pH condition produced more peptides with one or more missed cleavage sides. All three potential degradation hotspots in CDR2 are located within a single tryptic peptide (XXXXXXXXD_52b_GXXN_56_TXXXD_61_SXR).

Figure 3 shows the extracted ion chromatograms, the relative quantification, and the fragment-ion spectra confirming succinimide formation at the D52b-G53 motif. No deamidation or succinimide formation was detected at N56-T57 or D61-S62. Because the Asu peptide coelutes with the isoAsp peptide, the extent of water loss during chromatography and electrospray was estimated using samples digested at pH 7.5, and was very low at approximately 2 %. The higher T0 level of succinimide in the unconjugated sdAb compared with the sdAb-NODAGA conjugate may be attributed to partial succinimide hydrolysis during the conjugation reaction performed at pH 8. As expected, the HCD fragment-ion spectrum of the unmodified peptide (Figure 3D) shows a complete y-ion series, including those covering the “DG” sequence motif. In contrast, the corresponding Asu peptide lacks the y11 and y12 ions. Thus, succinimide formation can be unambiguously assigned to the D52b-G53 sequence motif rather than to the other potential degradation sites N56T57 or D61-S62.

**Figure 3.**
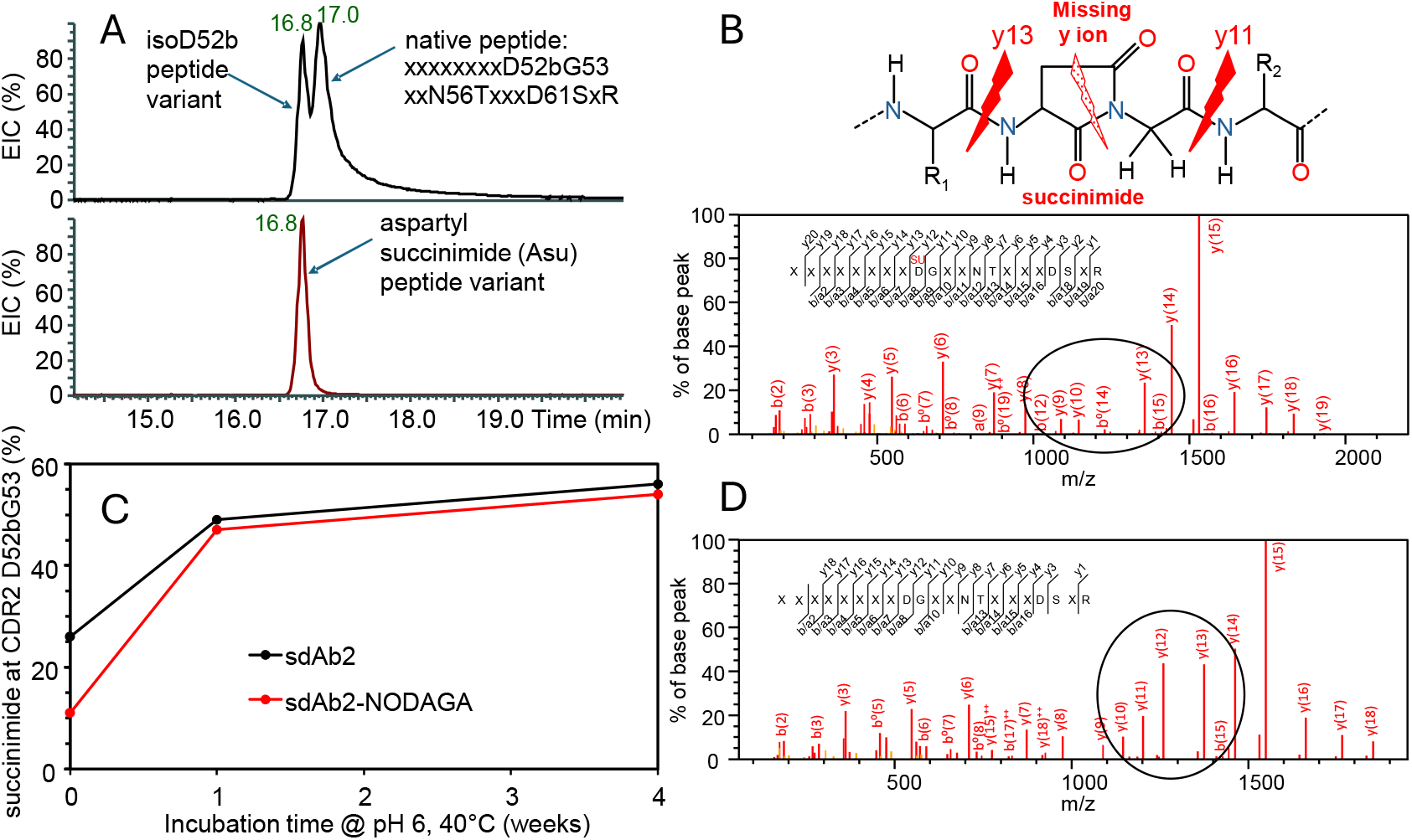
Confirmation of succinimide formation at sdAb2_D52b-G53 from tryptic peptide map at pH 6.0 by RP-C18-LC-MS/M analysis of sdAb and sdAb-chelator samples stored at pH 6 at 37 °C. A: Extracted ion chromatogram (EIC) of the tryptic peptide and the Asu variant of the peptide; B, D: fragment ion spectra of the Asu peptide variant (B) compared to the Asp peptide (D) and C: relative quantification of Asu formation at D52b-G53 after storage at pH 6 and 37 °C.

#### Isomerization of aspartate

can occur under both acidic and mildly basic conditions, particularly in sequence regions with high local flexibility, and may alter biological activity, increase susceptibility to proteolytic degradation, or even contribute to autoimmune responses. Isoaspartate analysis was performed at the tryptic-peptide level, where partial chromatographic resolution of the two peptide variants was achieved. Figure 4A illustrates that chromatographic separation was strongly dependent on the sample-preparation protocol: non-alkylated peptides generated using the trypsin digestion protocol at pH 6 showed improved peak resolution of the two isomers compared with the cysteine-alkylated peptide variants obtained from the digestion protocol at pH 7.5.

**Figure 4.**
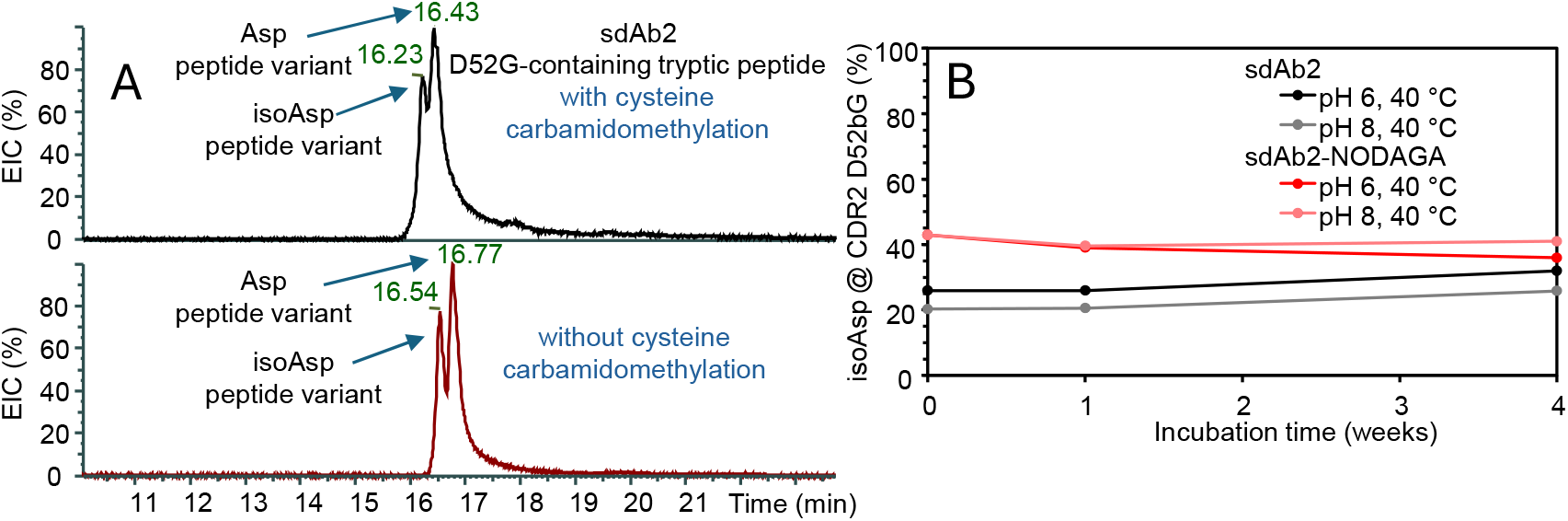
**A** Extracted ion chromatograms (EICs) of the tryptic peptide containing D52b-G53 and isoD52b-G53 in sdAb2 (exemplified for 4 weeks stress samples) using the digestion protocol at pH 6 without cysteine alkylation and at pH 7.5 with cysteine alkylation, respectively. **B** relative quantification of isoaspartate formation at D52b-G53 sequence motif.

Interestingly, the T_0_ samples exhibit a marked difference in the amount of isoAsp between the sdAb-chelator conjugate and the non-derivatized sdAb. This disparity is attributed to the alkaline pH conditions applied during the coupling reaction. Over the course of incubation, the difference gradually diminishes, and the isoAsp/Asp ratio stabilizes at approximately **1:2** for both tested pH conditions.

#### Fragmentation at the aspartate-proline (DP)

interface represents a well-known degradation motif, capable of undergoing hydrolysis under mildly acidic conditions and recognized as one of the most frequent degradation pathways in monoclonal antibodies [25]. It was evaluated using intact-mass RP-C4-LC-MS analysis. The shorter C-terminal hydrolysis product appears as a chromatographic pre-peak and is attributable to on-column hydrolysis at pH 2.4 and 75 °C during chromatography. Importantly, this peak does not increase in samples stored at 37 °C and pH 6 for either of the two sdAbs, as shown in Figure 5A and 5B. The larger N-terminal fragment of the hydrolyzed peptide could not be detected, as indicated by arrows in Figure 5C and D. Consequently, this potential liability is considered low-risk. When chromatography was performed at 50 °C, it further confirmed that hydrolysis at the DP motif was method-induced, rather than present in the stressed samples.

**Figure 5.**
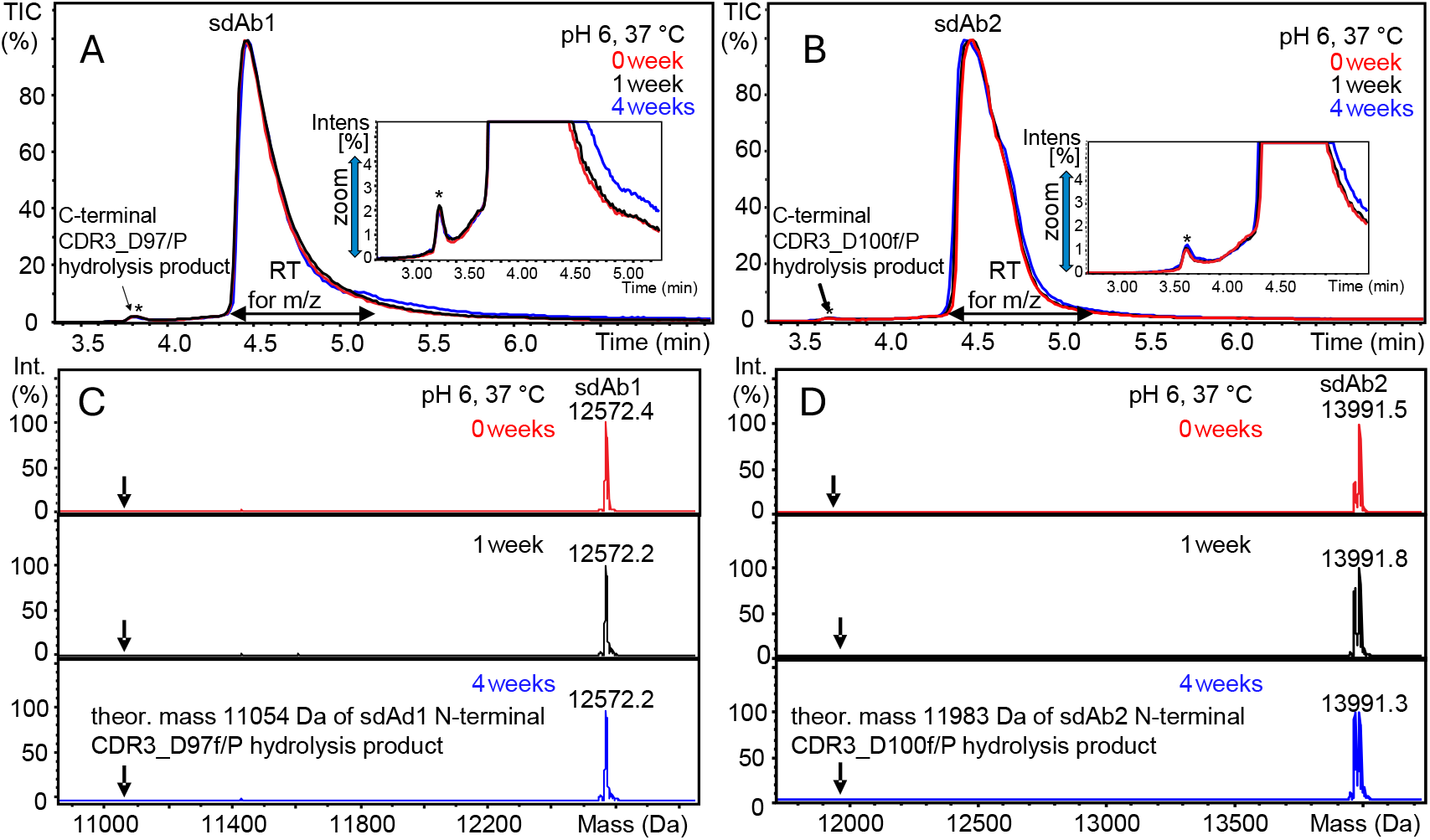
Assessment of hydrolysis tendency at DP sequence motif in CDR3 by intact mass RP-C4-LC-MS analysis of sdAb samples stored at pH 6 and 37 °C. A, B: total ion chromatograms and C, D charge-deconvoluted mass spectra of sdAb samples stored for different time periods. Spectra were obtained by summing up all spectra acquired across the main chromatographic peak as indicated by the double arrow in A and B.

### 3.4. Assessment of sdAbs Framework Stability and Stability of Chelator Conjugation

#### Pyroglutamate formation at the N-terminus

is of particular relevance because it prevents N-terminal amine-reactive conjugation. Relative quantification was performed based on intact mass analysis in cases where no additional water loss within the molecule was expected (e.g., absence of succinimide formation). In addition, relative quantification by tryptic peptide mapping yielded comparable results. Initial levels of N-terminal pyroglutamate formation ranged from 1.4 - 1.7% and increased to 2.4% following incubation at pH 6 and 40 °C for four weeks, and to 12% after incubation at pH 8. Both sdAbs exhibited similar trends (Figure S1).

#### Non-enzymatic hydrolysis

within the sdAb framework may impair chelator attachment, antigen-binding capacity, or overall molecular integrity. Fragmentation was first evaluated by intact LC-MS analysis. A progressively increasing chromatographic pre-peak corresponding to the chelator-free sdAb was observed in all sdAb-chelator conjugates. Fragmentation occurred at E1/V2, V2/Q3, and E6/S7 within the N-terminal region of the sdAbs and increased over time, with higher levels detected at pH 6.0 compared to pH 8.0 at 40 °C (*Figure 6*).

**Figure 6.**
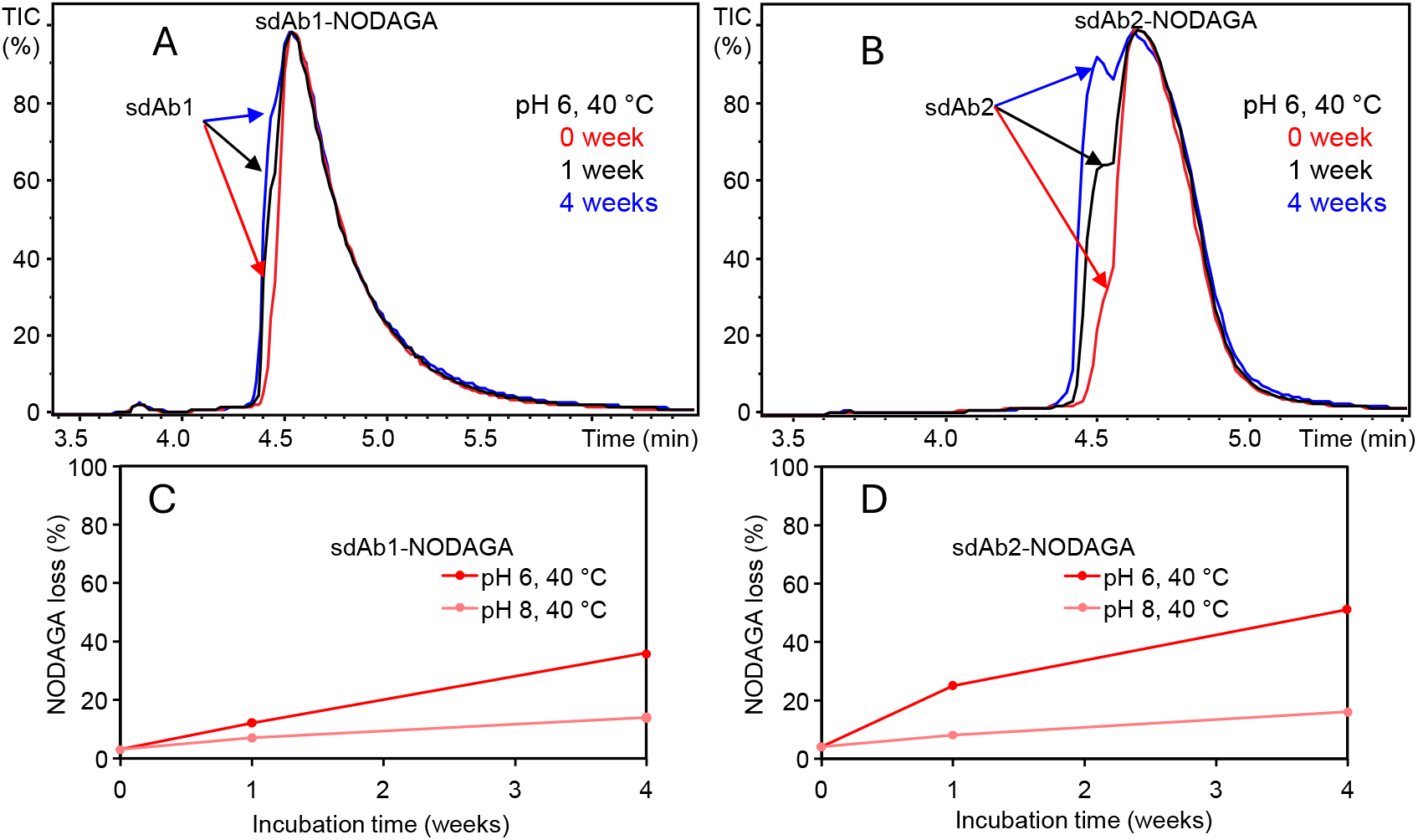
Assessment of loss of chelator by RP-C4-LC-MS analysis of intact sdAb samples stored at pH 6 and 8 at 40 °C. A-B: total ion chromatograms of sdAb molecules after storage at pH 6. C-D: relative quantification of chelator loss using peak intensities of charge-deconvoluted mass spectra.

In addition to chelator loss observed at both pH conditions, sdAb2 showed marked proteolytic degradation, evident as a pronounced chromatographic pre-peak in RP-C4-MS analysis of intact samples (*Figure 7*) and as an increased relative abundance of semitryptic peptides in the corresponding peptide maps (Figure S2). In contrast, sdAb1 remained stable when stored at pH 8 and elevated temperature.

**Figure 7.**
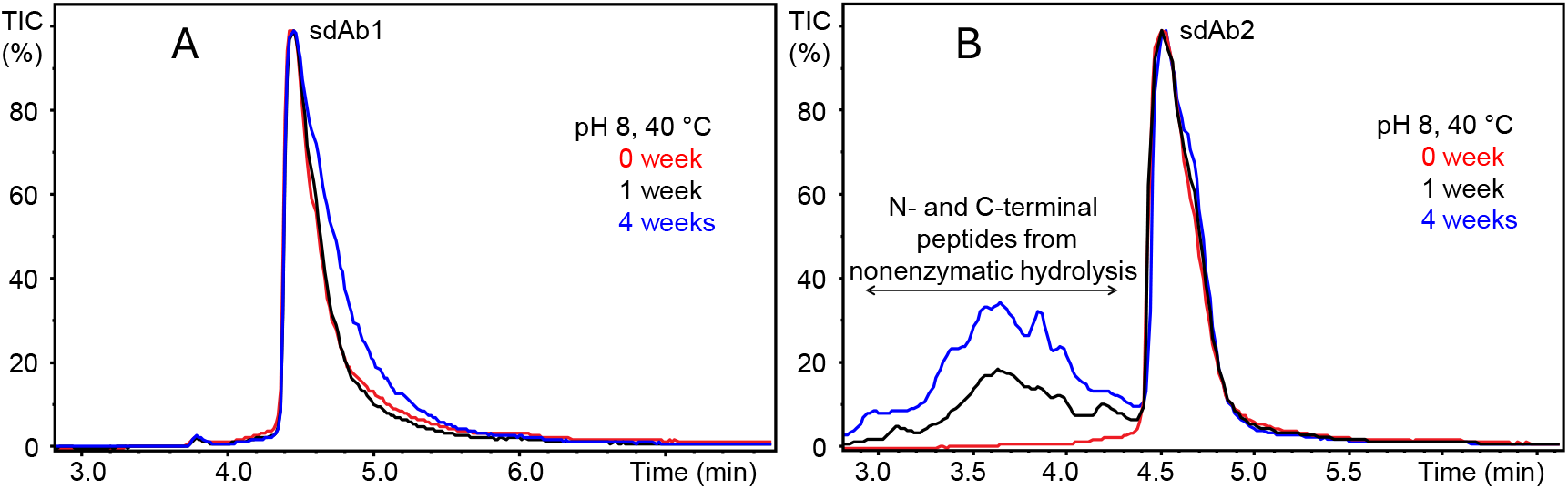
Assessment of proteolytic stability of sdAbs by RP-C4-LC-MS analysis of intact samples stored at pH 8 at 40 °C. A-B: total ion chromatograms of sdAb molecules after storage at pH 8.

### 3.5. Assessment of sdAbs Structural Integrity

The stability of the three-dimensional structure of the sdAbs was assessed by monitoring changes in intrinsic tryptophan fluorescence during thermal unfolding (*Figure 8*). sdAb1 contains three tryptophan residues. Based on the conserved VHH immunoglobulin fold and their positions within the sequence, we assume that one tryptophan is buried within the hydrophobic core formed by the β-sheet framework, whereas two residues are solvent-exposed. sdAb2 contains two tryptophans, one located in the hydrophobic core and one solvent-exposed.

**Figure 8.**
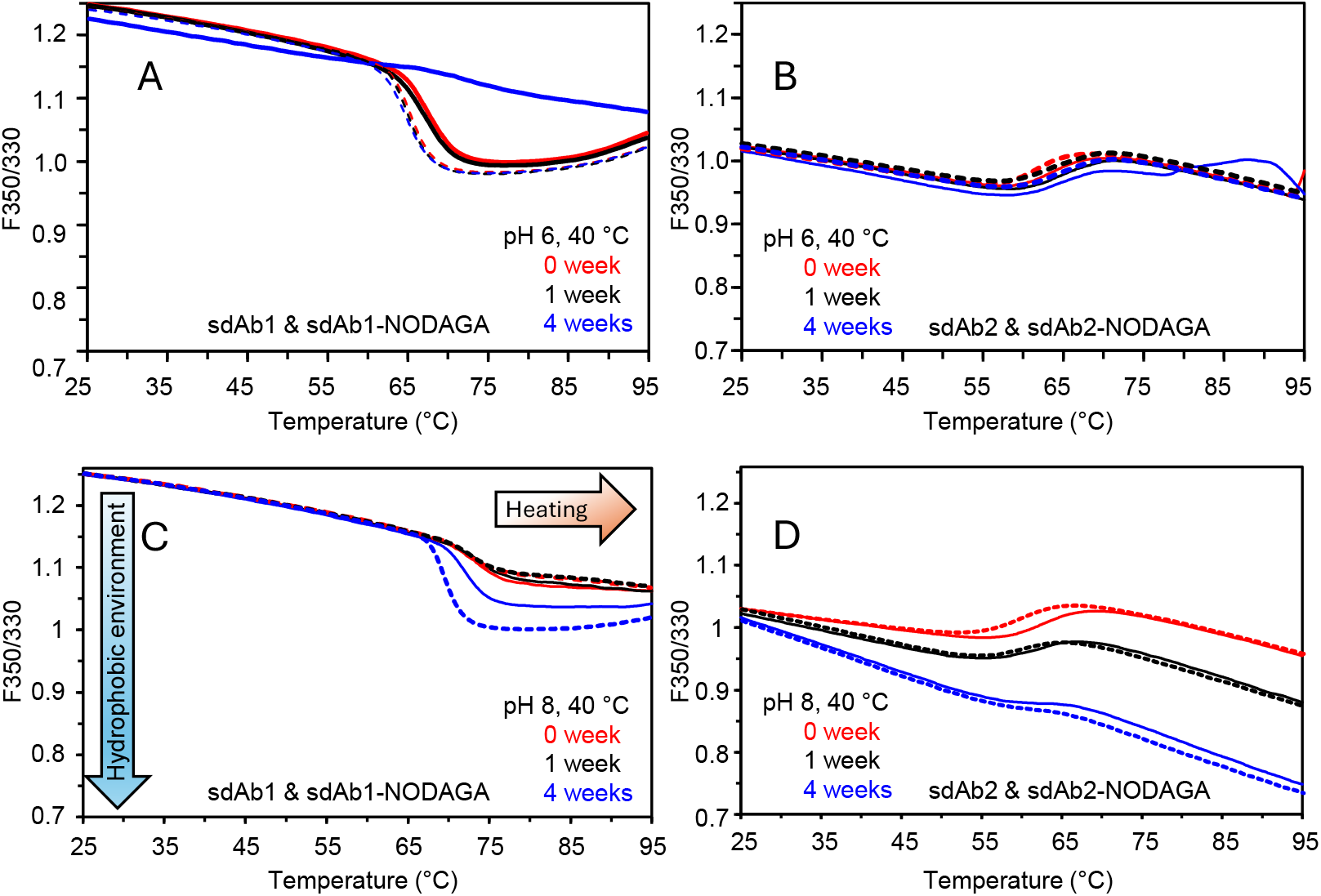
Thermal stability of sdAb samples (1 mg/mL) assessed by nanoDSF. The F350/330 tryptophan fluorescence ratio is shown for sdAb1 and sdAb2 after storage at pH 6 and 40 °C (A, B) and pH 8 and 40 °C (C, D). Solid lines represent unconjugated sdAbs; dashed lines represent NODAGA-conjugated sdAbs.

At 25 °C, sdAb1 exhibits a fluorescence ratio (F350/330) greater than 1, consistent with the presence of multiple solvent-exposed tryptophans. The ratio decreases progressively with increasing temperature. At approximately 60 °C, both sdAbs show a distinct fluorescence transition in samples stored at pH 6. At pH 8, the transition occurs at higher temperatures (> 65 °C) and appears less pronounced. In contrast, sdAb2 displays an F350/330 ratio of approximately 1 at 25 °C, consistent with fewer solvent-exposed tryptophan residues. Upon temperature increase, sdAb2 samples stored at pH 8 exhibit a rapid decrease in F350/330 values, indicating an enhanced contribution of hydrophobicity in the tryptophan environment under these conditions.

The fluorescence thermograms of unconjugated and NODAGA-conjugated sdAbs show similar unfolding profiles, with the exception of the sdAb1 samples stored for four weeks. Whereas the unconjugated sdAb1 sample exhibited no clear transition after four weeks at pH 6 and 40 °C, the corresponding NODAGA-conjugated sample retained a transition pattern comparable to earlier time points. This observation suggests a potential stabilizing effect of the NODAGA modification.

To further investigate the instability observed for sdAb2 and its NODAGA conjugate under basic pH conditions and elevated temperature, we analyzed the hydrodynamic radii and polydispersity indices (PDI) of sdAb2-NODAGA using DLS (*Figure 9*). To specifically capture changes in intrinsic molecular structure, only values corresponding to the main molecular population were considered.

**Figure 9.**
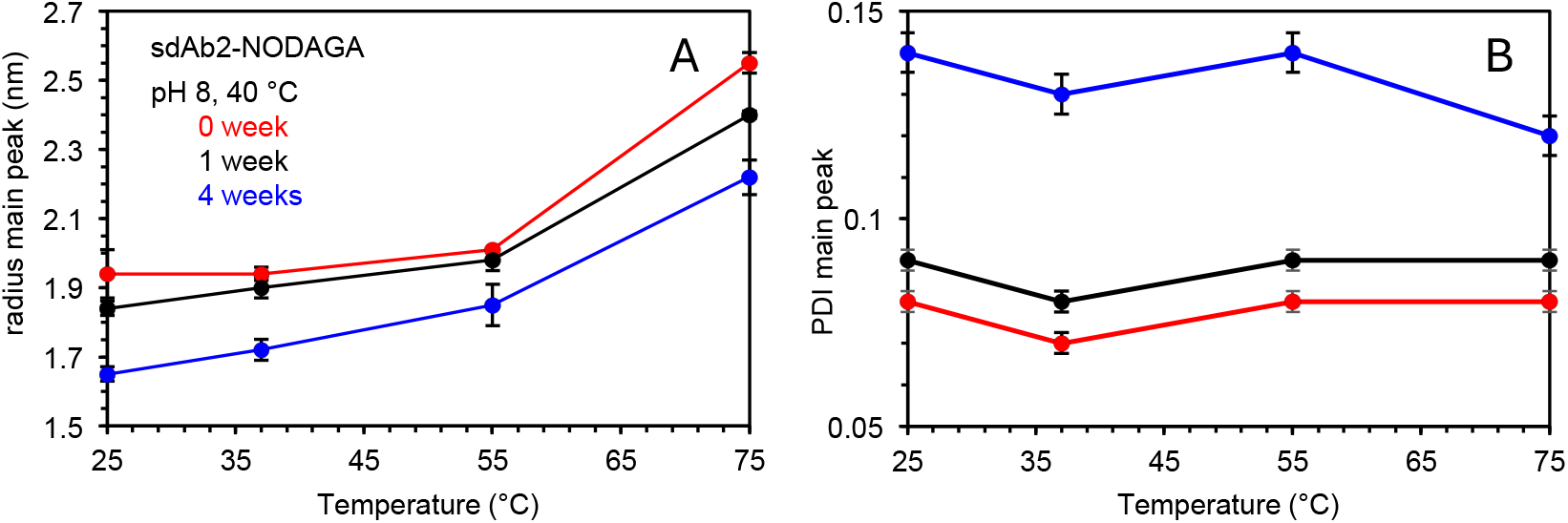
Hydrodynamic radius (A) and polydispersity index (PDI) (B) of sdAb2-NODAGA stability samples at pH 8, analyzed by DLS. Values refer to the main molecular population (main peak) and do not represent cumulant-based radius or PDI.

The data show that the main sdAb population decreases in hydrodynamic radius when stored at elevated temperature and pH 8. Concurrently, the polydispersity increases with storage time, indicating enhanced structural heterogeneity. However, the thermal unfolding behavior inferred from the radius and PDI thermograms remains unchanged. Similar observations were made for the unconjugated sdAb2 (data not shown).

### 3.6. Antigen Binding Capacity Assay by SEC-UV

A rapid and sensitive assay to evaluate the antigen-binding capacity of sdAbs and their conjugates is essential for in-process control and product release during manufacturing. The SEC-UV-based assay established here enables monitoring of sdAb stability with respect to antigen binding, as well as evaluating the antigen-binding properties after metal chelation. When combined with a radioactivity detector, the method also allows simultaneous assessment of radioactive payload integrity and target binding.

The underlying principle relies on the shift in elution time during size-exclusion chromatography (SEC) between the free sdAb and the sdAb-antigen complex (*Figure 10* B). Upon binding to the antigen, the sdAb peak disappears from its native elution position, while the antigen peak appears as a double signal: an early-eluting peak corresponding to the sdAb-antigen complex and a second peak corresponding to unbound antigen. A molar antigen-to-sdAb ratio of 2:1 was used to ensure complete binding of sdAb molecules. For reliable quantification, high-quality antigen at sufficient concentration and with an appropriate molecular weight difference relative to the sdAb is required. Commercial antigen formats proved insufficiently stable; therefore, an antigen-Fab construct was designed, expressed, and purified in-house. Specific sdAb binding to the antigen portion of the construct - rather than to the Fab domain - was confirmed.

**Figure 10.**
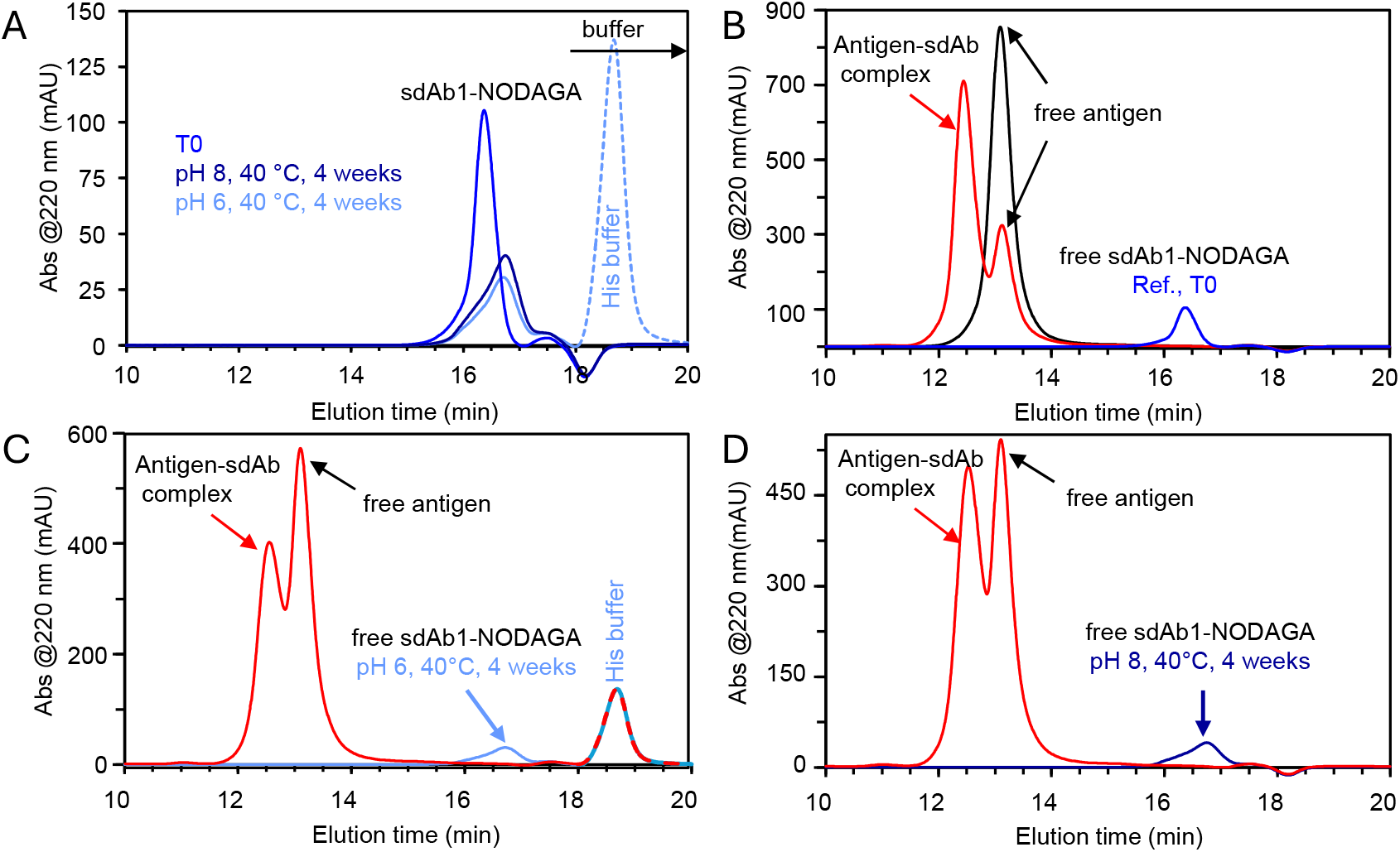
Antigen-binding capacity assessment by SEC-UV analysis. (A) Size-exclusion chromatogram of sdAb1-NODAGA samples. (B) Elution profiles of free sdAb1-NODAGA (blue), free antigen (black), and a 2:1 molar antigen:sdAb-NODAGA mixture (red). (C-D) Elution profiles of free sdAb1-NODAGA stability samples and corresponding sdAb–antigen complexes (∼1.5:1 antigen:sdAb molar ratio) for pH 6 and pH 8 stability conditions, respectively.

*Figure 10* shows that stability samples of sdAb1 exhibit peak shifts toward lower molecular-weight (LMW) species upon SEC analysis. This observation is consistent with the decreasing hydrodynamic radii observed at pH 8 and 40 °C (*Figure 9*A), resulting in peak broadening of the sdAb signal. Nevertheless, for both sdAb stability conditions the sdAb– NODAGA peak fully disappears upon incubation with the antigen (*Figure 10*C,D), while a new peak corresponding to the sdAb-antigen complex emerges. Thus, despite altered folding or size in the stability samples relative to the reference material, antigen binding of the sdAb is maintained.

### 3.6. Assessment of Chelator Ability to Bind Copper

Although no chemical degradation was detected in the chelator moiety of the conjugate, the copper-binding capability of the chelator was evaluated using mass spectrometry. For this purpose, an equimolar amount of copper chloride was added to sdAb1-NODAGA samples formulated in different buffer systems, followed by analysis via RP-C4-LC-MS. As shown in *Figure 11*, complex formation strongly depended on the copper-binding properties of the respective buffer. Quantitative Cu^2+^ chelation was observed in phosphate and TRIS buffer, whereas only approximately two-thirds of the copper was bound in histidine buffer. Consequently, assessment of copper-binding stability could not be performed for the pH 6 stability samples, as these had been prepared in histidine buffer.

**Figure 11.**
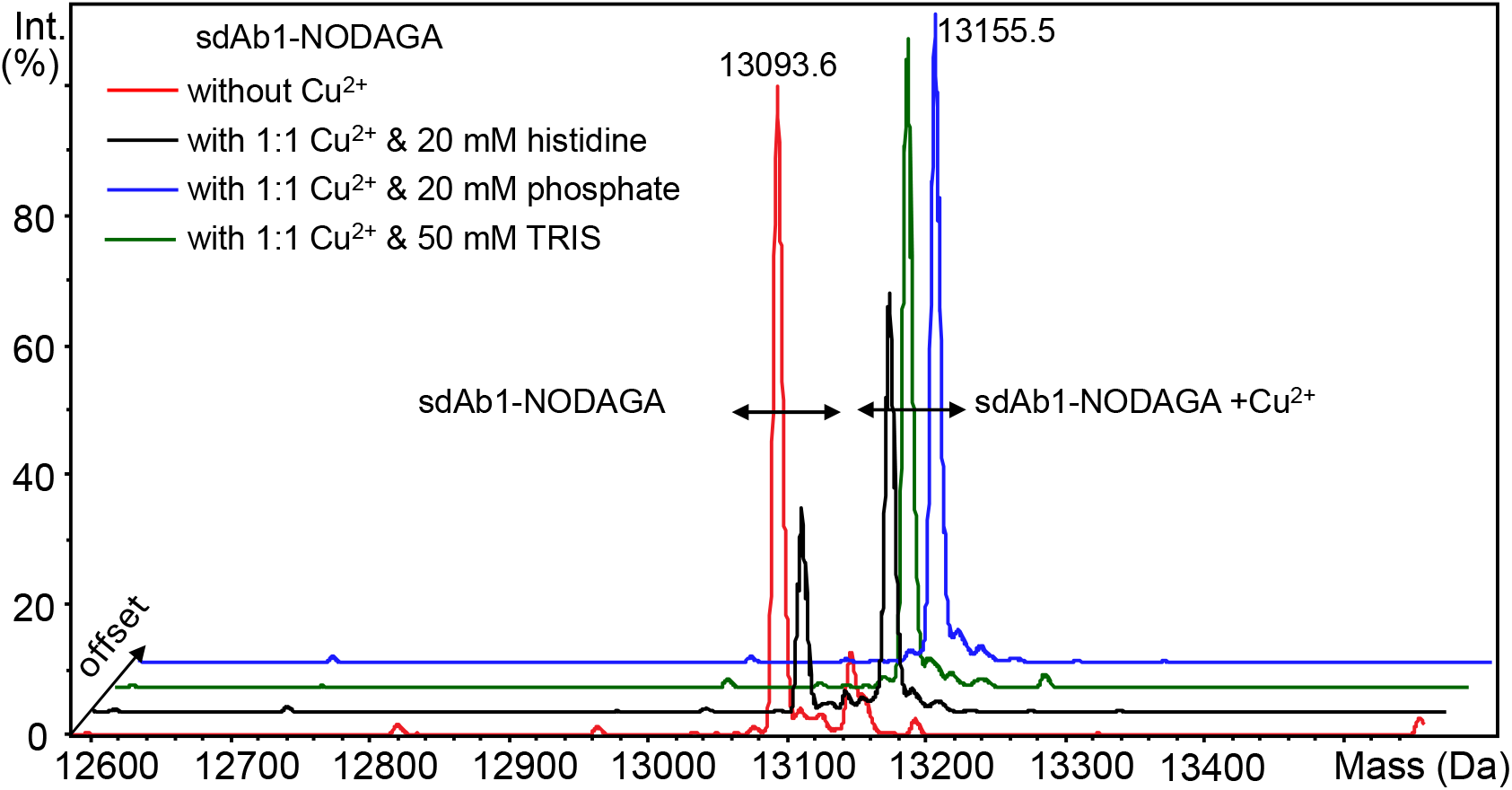
Overlay of mass spectra of sdAb-chelator conjugates incubated with Cu^2+^ in various buffer systems.

### 3.7. Assessment of Freeze-Thaw Stability

Repeated freeze-thaw cycles can induce structural perturbation or aggregation, depending on buffer composition and protein concentration. To evaluate freeze-thaw robustness, sdAb1 samples were subjected to 12 freeze-thaw cycles and subsequently analyzed by dynamic light scattering (DLS) and nanoDSF at a concentration of 1 mg/mL in formulation buffer (20 mM sodium phosphate, pH 6.8). As shown in *Figure 12*, no significant changes in hydrodynamic radius or PDI were detected after 12 cycles, indicating high structural stability of both sdAb1 and sdAb1-NODAGA. Thermal unfolding profiles likewise remained unchanged, confirming preservation of conformational stability.

**Figure 12.**
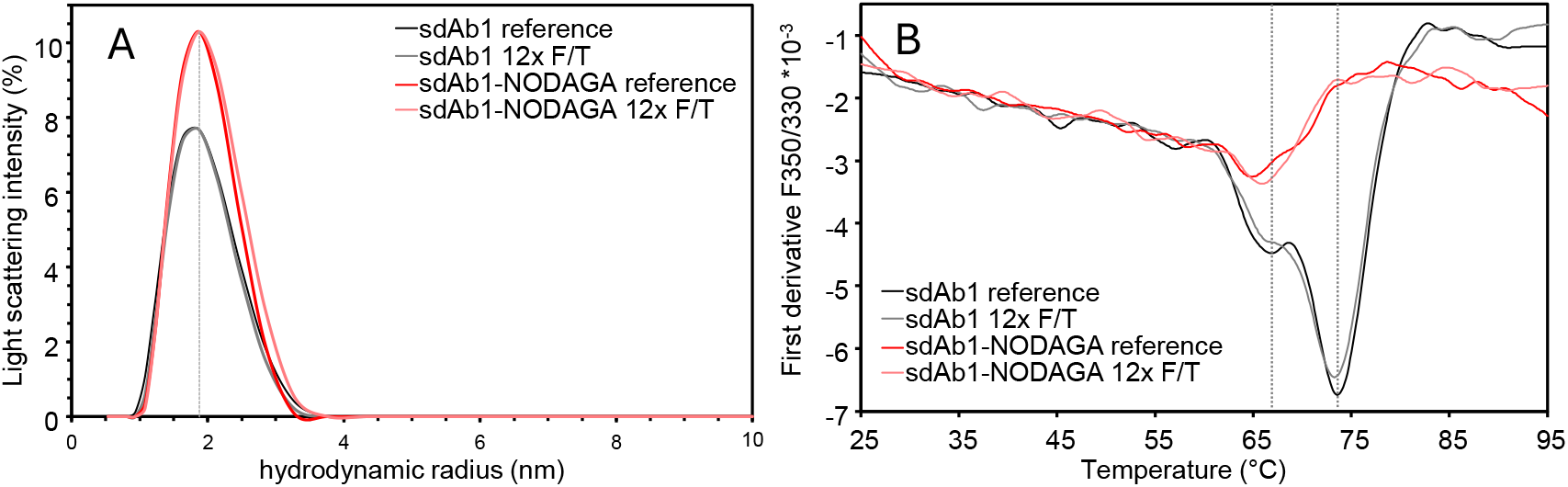
Assessment of structural stability of sdAb samples in formulation buffer (pH 6.8) after 12 freeze-thaw cycles by (A) DLS (hydrodynamic radius, PDI) and (B) nanoDSF (F350/330 ratio; three tryptophans in sdAb primary sequence).

### 3.8. Engineering the Asp-Gly Degradation Hotspot

K_D_ determination of the reference and the one-week stability sample did not reveal significant impact on binding affinity for sdAb2 even though the relative succinimide concentration increased by almost factor 2 (*Table 3*). This finding suggests that the CDR2 is not involved in the binding to the antigen. Nevertheless, knock-out of the degradation hot spot was attempted. Based on previous reports demonstrating successful mitigation of Asp isomerization hotspots [25], two sdAb variants were generated and evaluated for binding affinity. One variant contained a charge-conserving substitution of the Asp residue within the Asp-Gly hotspot to Glu, while the second variant replaced the glycine at the N+1 position with alanine. Compared to the wild-type (WT) sdAb2, the Glu-Gly variant displayed a seven-fold reduction in binding affinity, whereas the Asp-Ala variant showed only a two-fold decrease.

**Table 3.**
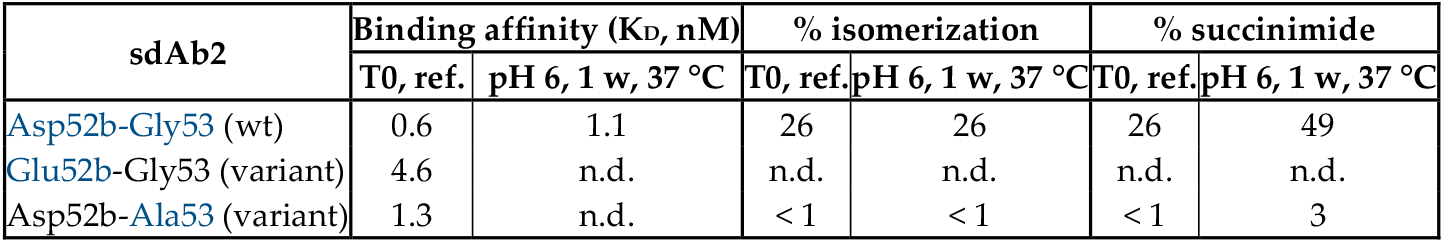
Binding affinity and chemical stability of sdAb2 variants with modifications in the Asp-Gly degradation hotspot.

| sdAb2 | Binding affinity ($K_D$, nM) | | % isomerization | | % succinimide | |
| --- | --- | --- | --- | --- | --- | --- |
|  | T0, ref. | pH 6, 1 w, 37 °C | T0, ref. | pH 6, 1 w, 37 °C | T0, ref. | pH 6, 1 w, 37 °C |
| Asp52b-Gly53 (wt) | 0.6 | 1.1 | 26 | 26 | 26 | 49 |
| Glu52b-Gly53 (variant) | 4.6 | n.d. | n.d. | n.d. | n.d. | n.d. |
| Asp52b-Ala53 (variant) | 1.3 | n.d. | < 1 | < 1 | < 1 | 3 |

As summarized in *Table 3*, neither isomerization nor succinimide formation at Asp52b was detected when glycine was substituted by alanine, while antigen-binding affinity remained largely preserved. A minor amount of succinimide (< 1% relative abundance) was observed in the one-week stress sample at pH 6 and 37 °C for the Asp-Ala variant, appearing as a chromatographically separated peak within the same tryptic peptide.

## 4. Discussion

The aim of this study was to establish a structured developability assessment for single-domain antibodies and their chelator conjugates. While sdAbs benefit from an intrinsically stable β-sheet scaffold and favorable biophysical properties, the results presented here show that their developability profile can be highly sequence-dependent and influenced by conjugation chemistry as well as environmental conditions during production and radiolabeling.

A central observation is the strong contrast between the two sdAb candidates tested. SdAb1 demonstrated a highly robust stability profile across all analytical readouts, with no measurable chemical degradation, structural destabilization, or loss of functional activity. In contrast, sdAb2 showed distinct liabilities concentrated within a single CDR motif, where succinimide formation and aspartate isomerization occurred rapidly. This highlights the importance of integrating intact-mass and peptide-level analytics to localize degradation pathways with high precision. The engineering experiments for sdAb2 show that targeted substitution of the glycine at the N+1 position of the Asp-Gly hotspot effectively eliminated succinimide formation and isomerization while preserving binding affinity, whereas the Asp→Glu substitution impaired affinity. This is consistent with known mechanisms of Asp-Gly instability and illustrates how minimal sequence adjustments can improve chemical robustness without extensive re-engineering.

The interplay between sequence context and conjugation chemistry further underscores the need for tailored assessment strategies. Succinimide abundance in sdAb2 decreased after NODAGA coupling, likely reflecting hydrolysis during the alkaline reaction. Although the chelators themselves remained chemically intact, copper-binding efficiency varied substantially between buffers, revealing a practical constraint for radiolabeling workflows. N-terminal fragmentation observed under accelerated conditions additionally highlights the importance of formulation pH and buffer selection when the N-terminus serves as a conjugation site.

While SEC-based workflows have been used to identify functionally relevant chemical modifications in IgGs - such as competitive separation of modified and unmodified antibody species [26] and paratope-level mapping [27] - these approaches cannot be directly transferred to sdAbs because of their much smaller molecular weight (∼14 kDa). SdAbs elute near the lower resolution limit of standard SEC stationary phases, requiring high-resolution columns with smaller pore sizes and an antigen size chosen to yield distinct peaks for free sdAb, free antigen, and the sdAb-antigen complex.

These constraints make sdAb-specific SEC configurations essential for reliably detecting binding losses and functional liabilities.

Radiolabeling can also alter protein integrity and antigen binding, as shown by experimental studies demonstrating radiolysis-driven structural damage and reduced binding potency in labeled antibodies [28]. Given that nanobodies are particularly sensitive to such effects due to their small size, radiolabeling conditions must be carefully optimized [29]. In this context, the sdAb-adapted SEC-binding-ability assay established here provides an essential and radiochemistry-compatible, high-resolution tool to ensure that radiolabeling does not impair antigen binding.

Collectively, the analytical workflow established in this study demonstrates that sdAbs exhibit unique degradation behaviors compared to conventional antibodies and therefore require dedicated developability strategies. The stepwise, hierarchical approach used here minimizes experimental burden while enabling sensitive detection of critical liabilities. Importantly, the combination of mass spectrometry, structural analytics, and functional assays provides a comprehensive framework that can be readily adapted to other sdAbs or conjugation chemistries used in diagnostic or therapeutic applications.

## 5. Conclusions

This study presents an integrated and efficient approach for the developability assessment of single-domain antibodies and their chelator conjugates. The strategy successfully identified molecule-specific degradation hotspots, structural vulnerabilities, and buffer-dependent behaviors relevant for long-term storage and radiolabeling. While one sdAb candidate exhibited high intrinsic stability, the second displayed a pronounced CDR-localized instability that was effectively mitigated through targeted protein engineering. The combined use of mass spectrometry, structural assays, and functional binding analytics provides a practical and broadly applicable framework that supports candidate selection, formulation development, and process optimization for sdAb-based radiopharmaceuticals. Overall, the results highlight the necessity of sdAb-adapted analytical workflows and demonstrate how early developability testing can significantly de-risk downstream manufacturing and clinical translation.

## Supplementary Materials

available.

## Author Contributions

Conceptualization, Ph.K. and B.T. and A.Z.; methodology, Ph.K. and A.Z.; analysis, S.M., E.H., S.S.; data curation, A.Z.; writing-original draft preparation, A.Z.; writing—review and editing, Ph.K., B.T.; All authors have read and agreed to the published version of the manuscript.

## Data Availability Statement

The data that support the findings of this study are available from the corresponding authors upon reasonable request.

## Abbreviations

AGC: Automatic Gain Control
Asu: Succinimide intermediate (aspartimide)
BSA: bovine serum albumine
BLI: Biolayer Interferometry
CDR: Complementarity-Determining Region
CID: Collision-Induced Dissociation
EIC: extracted ion current
ESI: electrospray ionization
DLS: Dynamic Light Scattering
DMSO: Dimethyl Sulfoxide
DPBS: Dulbecco’s Phosphate-Buffered Saline
DTT: Dithiothreitol
ESI: Electrospray Ionization
Fab: Fragment antigen-binding
Gua-HCl: Guanidine Hydrochloride
HCD: Higher-Energy Collisional Dissociation
HPLC: High-Performance Liquid Chromatography
K_D_: equilibrium dissociation constant
LC–MS: Liquid Chromatography-Mass Spectrometry
LC-MS/MS: Liquid Chromatography-Tandem Mass Spectrometry
LMW: Low Molecular Weight
mAU: milli-Absorbance Units
MWCO: Molecular Weight Cut-Off
nanoDSF: Nano Differential Scanning Fluorimetry
NCS: Isothiocyanate functional group (–N=C=S)
n.d.: not determined
NODAGA: Asu1,4,7-Triazacyclononane-1,4,7-triacetic acid-glutaric acid
PBS: phosphate-buffered saline
PDI: Polydispersity Index
PET: Positron Emission Tomography
QTOF: Quadrupole Time-of-Flight Mass Spectrometer
RP-C4 / RP-C18: Reversed-Phase Chromatography with C4/C18 stationary phase
RPM: Revolutions Per Minute
RT: Room Temperature
SA biosensor: Streptavidin biosensor
sdAb: Single-Domain Antibody
SEC: Size-Exclusion Chromatography
TCEP: Tris(2-carboxyethyl)phosphine
TFA: Trifluoroacetic Acid
TIC: total ion current
UV: Ultraviolet Light Detection
VWD: variable wavelength detector
w: week
wt: wild type

