## Supplemental Figures for "Developability Evaluation of Single-Domain Antibody Chelator Conjugates for Diagnostic Radiotracers"

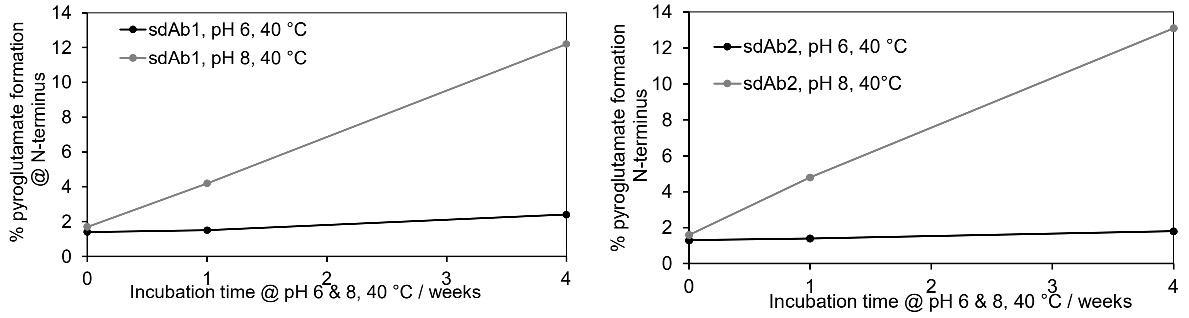


Figure S1: Pyroglutamate formation at N-terminal glutamic acid as relatively quantified from the N-terminal tryptic peptide in LC-MS/MS analysis.

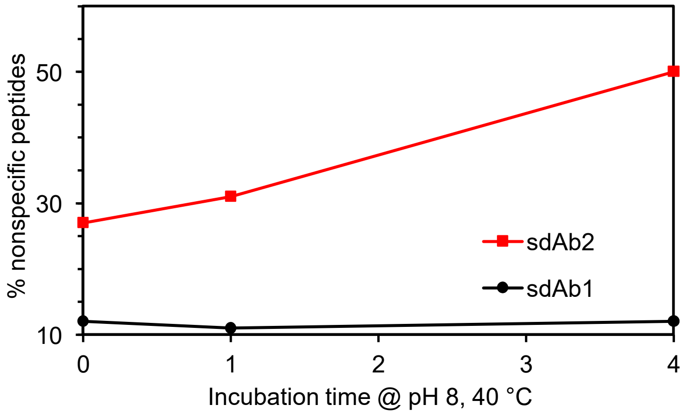


Figure S2: Relative amount of nontryptic peptides observed after tryptic digestion and peptide map analysis. The peak areas of EICs of nontryptic peptides was divided by the total peak area of all assigned peaks in LC-MS/MS analysis.

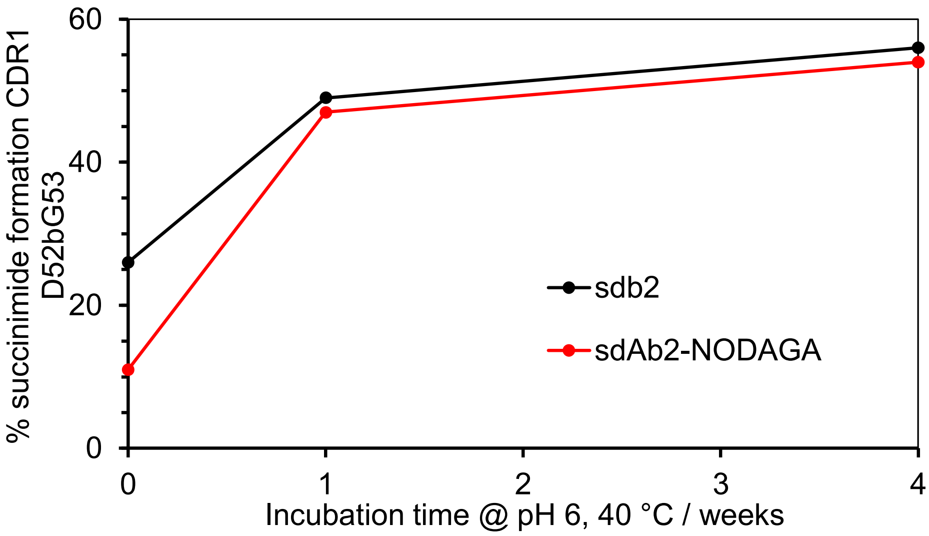


Figure S3: Relative amount of succinimide found in sdAb2 at D52bG53 as quantified from the tryptic peptide by LC-MS/MS analysis
